# Inactivation of CDK4/6, CDK2, and ERK in G1-phase triggers differentiation commitment

**DOI:** 10.1101/2025.04.07.647597

**Authors:** Sanjeev Sharma, Henri Berger, Tobias Meyer, Mary N. Teruel

## Abstract

Terminal cell differentiation, a process vital for tissue development and regeneration where progenitor cells acquire specialized functions and permanently exit the cell cycle, is still poorly understood at the molecular level. Using live-cell imaging and adipogenesis as a model, we demonstrate that the initial stage involves a variable number of cell divisions driven by redundant CDK4/6 or CDK2 activation.. Subsequently, a delayed decrease in cyclin D1 and an increase in p27 levels leads to the attenuation of CDK4/6 and CDK2 activity. This results in G1 lengthening and the induction of PPARG, the master regulator of adipogenesis. PPARG then induces p21, and later p18, culminating in the irreversible inactivation of CDK4/6 and CDK2, and thus, permanent cell cycle exit. However, contrary to expectation, CDK inactivation alone is not sufficient to trigger commitment to differentiation and functional specialization; ERK inactivation is also required. Our study establishes that the coordinated activation and subsequent delayed inactivation of CDK4/6, CDK2, and ERK are crucial determinants for irreversible cell cycle exit and differentiation commitment in terminal cell differentiation.

## INTRODUCTION

The process of terminal cell differentiation produces specialized cell types, including neurons, muscle cells, and adipocytes (fat cells), which are essential for the proper functioning of tissues and organs^1–3^. This process generally begins with progenitor cell division, leading to an expansion o the progenitor pool, and culminates in permanent exit from the cell cycle and the development of specialized functions^1–3^. Failure of terminally differentiated cells to permanently exit the cell cycle is a defining feature of cancer^2,4^. Therefore, elucidating the mechanisms governing the timing of progenitor cell cycle exit and its connection to differentiation commitment is of fundamental importance.

When individual cells decide to stop proliferating and differentiate is highly variable and thus requires that the process be studied live in single cells^5^. We had previously developed such a method to investigate the terminal cell differentiation of adipocytes (fat cells)^5,6^. These studies revealed that preadipocytes commit to differentiate during the G1 phase of the cell cycle, but only if the expression of PPARG, the master transcriptional driver of adipogenesis, surpasses a critical threshold level (“differentiation commitment point”) at which multiple positive feedbacks keep PPARG levels persistently high. While these studies demonstrated that permanent cell cycle exit and differentiation commitment both occur during G1 phase, it was not known which molecular components control progenitor exit from the cell cycle exit and in particular, whether cell cycle exit precedes and is sufficient to trigger differentiation commitment.

In mammals, cell cycle entry and exit are controlled in G1 phase by the activity of two kinases, CDK4/6 and CDK2^7^. However, the timing, mechanisms, and necessity of these kinase activities in controlling cell cycle exit during terminal cell differentiation had not been previously investigated. Moreover, the distinct roles of CDK4/6 and CDK2 in cell cycle entry and exit in other cell types^7^ raised the question of whether both kinase activities are needed to regulate cell cycle exit during terminal differentiation. Furthermore, genetic deletion and overexpression studies showed that cyclin D1 and the three CDK inhibitors, p21, p27, and p18, which regulate CDK4/6 and CDK2, also have critical roles in regulating adipogenesis and other differentiation processes^8–10^. However, if and how these critical CDK regulators inactivate CDK4/6 and CDK2 during terminal cell differentiation was also not known.

To understand how these critical kinases are regulated, we developed a live, single-cell method based on mosaic reporter transfection to track individual cells while measuring changes in critical signaling activities that govern both proliferation and differentiation. Specifically, we simultaneously measured the activity of the CDK4/6 and CDK2 kinases, which control cell cycle entry, along with endogenous level of the master transcriptional regulator of adipogenesis, PPARG.

Markedly, we found that during terminal cell differentiation, CDK4/6 and CDK2 were inactivated after each mitosis and then either jointly increased to start another cell cycle or jointly remained inactive. Using recently-developed specific inhibitors for CDK2 and CDK4/6^11,12^, we demonstrated that either CDK4/6 or CDK2 alone could trigger cell cycle entry, but both must be inactivated to increase PPARG levels to the threshold required for differentiation commitment. We identified a sequential order for CDK4/6 and CDK2 inactivation preceding differentiation commitment: the process begins with a decrease in cyclin D1 and an increase of cyclin-CDK inhibitor p27, which together reduce CDK4/6 and CDK2 activity to lengthen G1 phase. The lengthening of G1 phase provides sufficient time for PPARG levels to reach the differentiation commitment threshold and drive expression of the cyclin-CDK inhibitor p21, and later the CDK4/6 inhibitor p18, to permanently suppress CDK2 and CDK4/6 activity. Unexpectedly, inactivation of both CDK4/6 and CDK2 was insufficient for increasing PPARG levels and a subsequent inactivation of ERK kinase was also needed. Together, our study reveals an ordered signaling process that ends a variable cell division period and drives progenitor differentiation based on the inactivation of CDK4/6 and CDK2 followed by inactivation of ERK.

## RESULTS

### Either CDK4/6 or CDK2 must be active in G1 to drive adipocyte progenitor proliferation

We investigated the relationship between proliferation and terminal cell differentiation in OP9 preadipocytes in which CRISPR-mediated genome editing had been used to tag endogenous PPARG with mCitrine(YFP) and a FUCCI cell cycle phase reporter (APC/C-mCherry) had been co-expressed^5,6^ (Figure 1A-D). First, we validated that addition of an adipogenic stimulus (DMI) causes PPARG levels to increase after an initial variable delay and that progenitor cells proliferate early in the differentiation process (Figure 1A, B)^5,6^. The DMI cocktail consists of the glucocorticoid analog dexamethasone, phosphodiesterase inhibitor IBMX to increase cAMP, and insulin^5^. We next validated that removal of the adipogenic stimulus after 2 days can be used to define a precise level of PPARG at which cells commit to differentiate, as shown by cells transitioning from a low PPARG (undifferentiated) state into a high PPARG (differentiated) state if they crossed the PPARG threshold level^5,6^ (Figure 1B).

**Figure 1.**
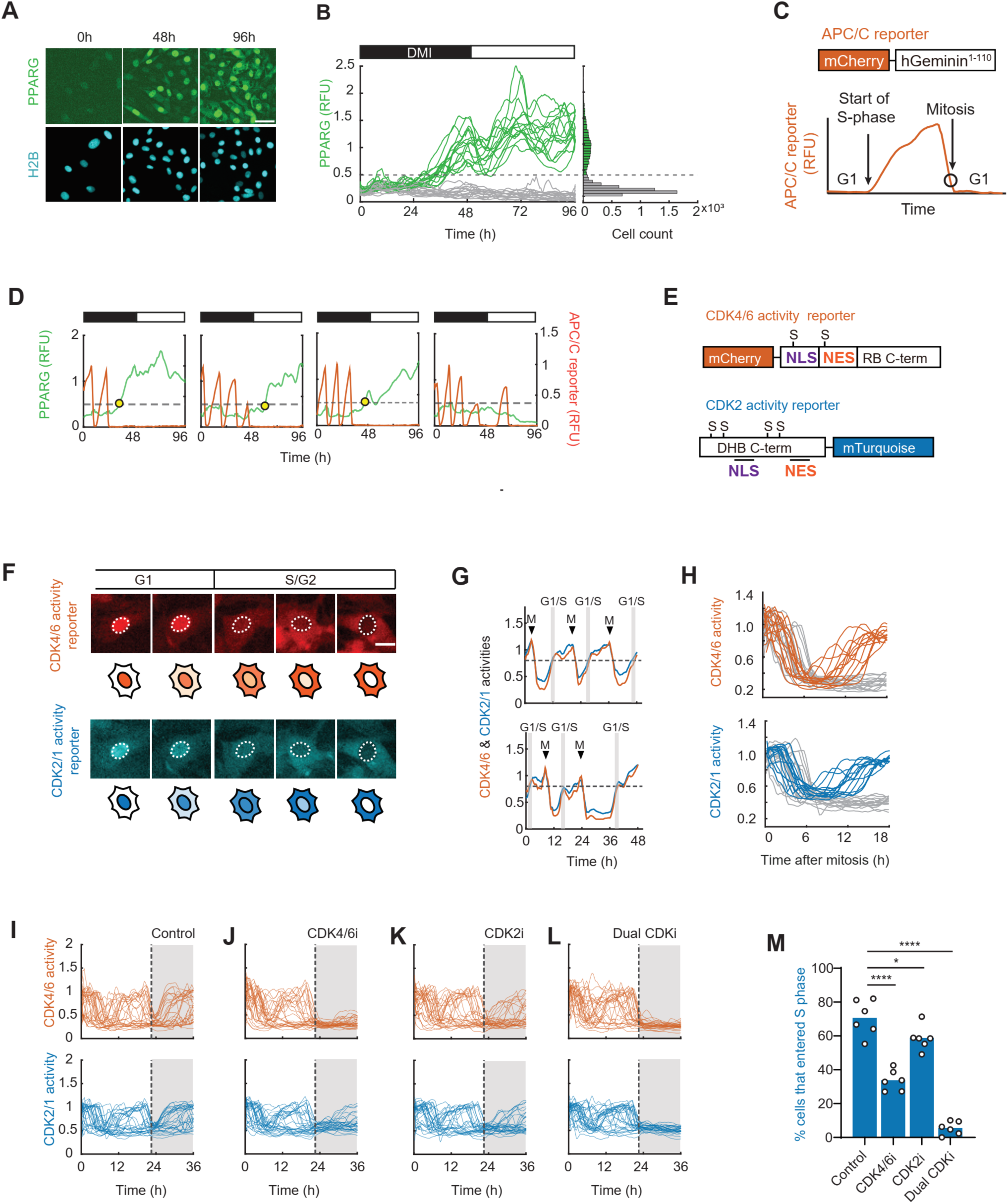
Adipocyte progenitor proliferation can be driven by CDK4/6 or CDK2 activation. (A) OP9 cells expressing mCitrine-PPARG and a nuclear marker (H2B-mTurquoise2) were differentiated over 4 days with the standard adipogenic DMI cocktail (Dexamethasone - 1 μM, IBMX – 250 μM, and Insulin – 1.75 nM). Scale bar - 50 μm. (B) Representative single-cell time courses of mCitrine-PPARG from a typical experiment. In the standard DMI-induced differentiation protocol, DMI cocktail is added to cell culture media for 48h (horizontal black bar), and media is replaced after 48h with fresh media containing just insulin for another 48h (white horizontal bar). 50 single-cell time courses are shown as examples. The differentiation commitment threshold is indicated by the dashed line. Cells go on to either differentiate by increasing PPARG levels above the threshold (green time courses) or remain undifferentiated with low PPARG levels (grey time courses). Distribution of PPARG levels at 96h shown on the right. Data shown is representative of >2000 cells tracked in the experiment and three biological replicates. (C) Schematic of APC/C cell cycle reporter construct and dynamics of the reporter’s nuclear fluorescence intensity within a cell during different phases of the cell cycle. (D) Dual reporter cells expressing mCitrine-PPARG and mCherry-APC/C-reporter were differentiated using the standard DMI protocol. Representative time courses from 4 single cells. The APC/C-reporter signal is low during the G1-phase, increases during the S/G2 phases and drops off at mitosis. In differentiating cells, commitment to differentiation (yellow circle) occurs during the G1-phase, whereas undifferentiated cells continue to divide intermittently during the 96h experimental window (bottom right panel). Data shown representative of >2000 cells tracked in the experiment and three biological replicates. (E) CDK4/6 and CDK2 kinase-translocation reporter (KTR) constructs. NLS - Nuclear localization signal sequence, NES – Nuclear export signal sequence, S – consensus serine phosphorylation site for CDKs. (F) Representative cells expressing both the CDK4/6 and CDK2 reporters. Increasing CDK activities during the cell cycle leads to the phosphorylation and translocation of the two reporters from the nucleus to the cytosol. Scale bar - 10 μm. (G) Single-cell time courses from two proliferating OP9 progenitor cells expressing CDK4/6 and CDK2 reporters. CDK4/6 and CDK2 activities were measured as the ratio of cytosolic to nuclear signal intensity. ‘M’ marks mitosis events. ‘G1/S’ marks the transition from G1 to S-phase, defined as when CDK2 and CDK4/6 reporter activities reach a value of 0.8, as indicated by the dashed line. (H) Single-cell timecourses of CDK4/6 (top) and CDK2 (bottom) activity from 30 OP9 progenitor cells that undergo mitosis (seen by the drop in CDK activities) within 6h of starting the imaging. CDK activity bifurcates into high (orange and blue) and low states (grey) after mitosis. Representative of two biological replicates. (I-L) DMSO (control), CDK4/6 inhibitor (CDK4i, Palbociclib, 1u M), CDK2 inhibitor (CDK2i, Tagtociclib, 1uM), or both CDK4/6 and CDK2 inhibitors (Dual CDKi, 1uM of each) were added to the culture media of proliferating OP9 progenitor cells 24h after the start of imaging (dashed vertical line). Plots show single-cell time courses from 30 cells that had undergone at least one mitosis event (CDK2 activity >1) since the start of imaging and were in G1-phase when drugs were added. Datapoints represent independent wells from the same experiment with over 100 cells analyzed per well. (M) Percent of cells from Figure 1I-L that entered S phase, as defined by the CDK2 signal increasing above a value of 0.8 within 16h of drug addition (see Methods).

We next measured PPARG levels together with the APC/C-mCherry cell cycle reporter, which drops sharply low at the end of mitosis and only rises again at the start of S/G2 phases^13,14^. G1-phase can thus be defined as the low period right after the sharp drop in the reporter (Figure 1C). We confirmed that cells only reached the threshold to commit to differentiation during G1-phase^5^ (Figure 1D). Furthermore, we observed that addition of the differentiation stimulus resulted in a variable number of cell divisions that only continued if PPARG levels remained below the threshold. Upon reaching the threshold, cells ceased cycling and exhibited a further increase in PPARG levels as they differentiated into mature adipocytes (Figure 1D). In contrast, cells that maintained PPARG levels below the threshold continued to divide, indicating their undifferentiated progenitor state and preservation of their proliferative potential (Figure 1D).

To elucidate the mechanisms controlling cell cycle exit during G1 phase, we focused on the two primary kinases regulating this phase: CDK4/6 and CDK2^7^. We engineered a triple-reporter cell line by stably introducing fluorescence reporters for CDK4/6 and CDK2/1 activities into OP9 preadipocyte cells expressing mCitrine(YFP)-PPARG (Figure 1E). These reporters undergo selective phosphorylation by CDK4/6 and CDK2/1, resulting in increased nuclear export as their respective CDK activities rise during G1 phase^15,16^ (Figure 1F). Consequently, CDK4/6 and CDK2/1 activities can be quantified by measuring the ratio of nuclear to cytoplasmic localization. Here, we refer to the CDK2/1 reporter as a CDK2 reporter, as CDK1 is generally inactive during G1 phase. Finally, we employed mosaic mixtures of OP9 preadipocytes with and without reporters to be able to effectively monitor individual fluorescently-labeled preadipocytes over several days (Figure 1F, also see Supplementary Figure 1 and Methods).

First, we measured CDK4/6 and CDK2 activities in cycling preadipocytes. Previous studies in epithelial cells had shown differences in the time courses of the two kinase activities, with CDK4/6 activity often staying high during and after mitosis despite CDK2 activity being low^17^. In contrast, we found that in preadipocytes, the CDK4/6 and CDK2 activities were closely correlated (Figure 1G). Following a drop in both activities after mitosis, both activities then jointly increase after variable delays, or both stay permanently low (Figure 1G and 1H). In most cell types, CDK4/6 activity is necessary for proliferation^7,18^, and addition of the specific CDK4/6 inhibitor Palbociclib^11^ strongly suppresses proliferation, as marked by the suppression of CDK2 reporter activity^17^. Here in OP9 preadipocyte cells, we also observed that Palbociclib strongly suppressed CDK4/6 activity. However, in many cells, CDK2 remained active, and proliferation continued (Figure 1I and J).

We next inhibited CDK2 activity using the recently developed selective small molecule inhibitor of CDK2, Tagtociclib (PF-07104091)^12^. We observed that cells still entered the cell cycle, but only after a delay as shown by the slower increase in the CDK2 reporter signal in Tagtociclib-treated cells compared to control cells (Figure 1I and 1K). It should be noted that we observed an increase in the CDK2 reporter signal despite addition of Tagtociclib. This increase in CDK2/1 reporter signal reflects a previously-identified mechanism in cells with inactive CDK2, where CDK1 activity increases after a delay, bypassing the need for CDK2 activity to start S-phase^19^. Consistent with CDK1 activation requiring prior CDK4/6 activation, adding Palbociclib and Tagtociclib together, completely prevented an increase in the CDK2 reporter signal (Figure 1L). A quantitative analysis further shows that the percent of cells that enter S-phase (i.e. proliferating cells) is completely suppressed only when both CDK4/6 and CDK2 are inhibited (Figure 1M).

We conclude the undifferentiated preadipocytes can proliferate by activating either CDK4/6 or CDK2 during G1 phase. Furthermore, they can only exit the cell cycle if both CDK4/6 and CDK2 are inactivated.

### Progenitor cells permanently exit the cell cycle during terminal differentiation by inactivating CDK4/6 and CDK2 in G1 phase

Having established the roles of CDK4/6 and CDK2 in unstimulated progenitor cells, we next measured the activity changes of these kinases in progenitor cells induced to differentiate. Upon adding the DMI adipogenic stimulus to the triple-reporter cells, we observed that CDK4/6 and CDK2 activities both became inactive after each mitosis. Subsequently, they either both increased after variable delays or remained persistently low (Figure 2A). Notably, when we added either a CDK4/6 or CDK2 inhibitor along with DMI, cells could still increase the CDK2 reporter signal after mitosis, albeit with a delay (Figure 2B, C). Complete suppression of proliferation during terminal differentiation was only achieved by adding both inhibitors (Figure 2D-E). Thus, consistent with our findings in unstimulated progenitor cells (Figure 1I-M), both CDK4/6 and CDK2 must be inactivated to terminate the proliferative period during terminal differentiation (Figure 2A-E).

**Figure 2.**
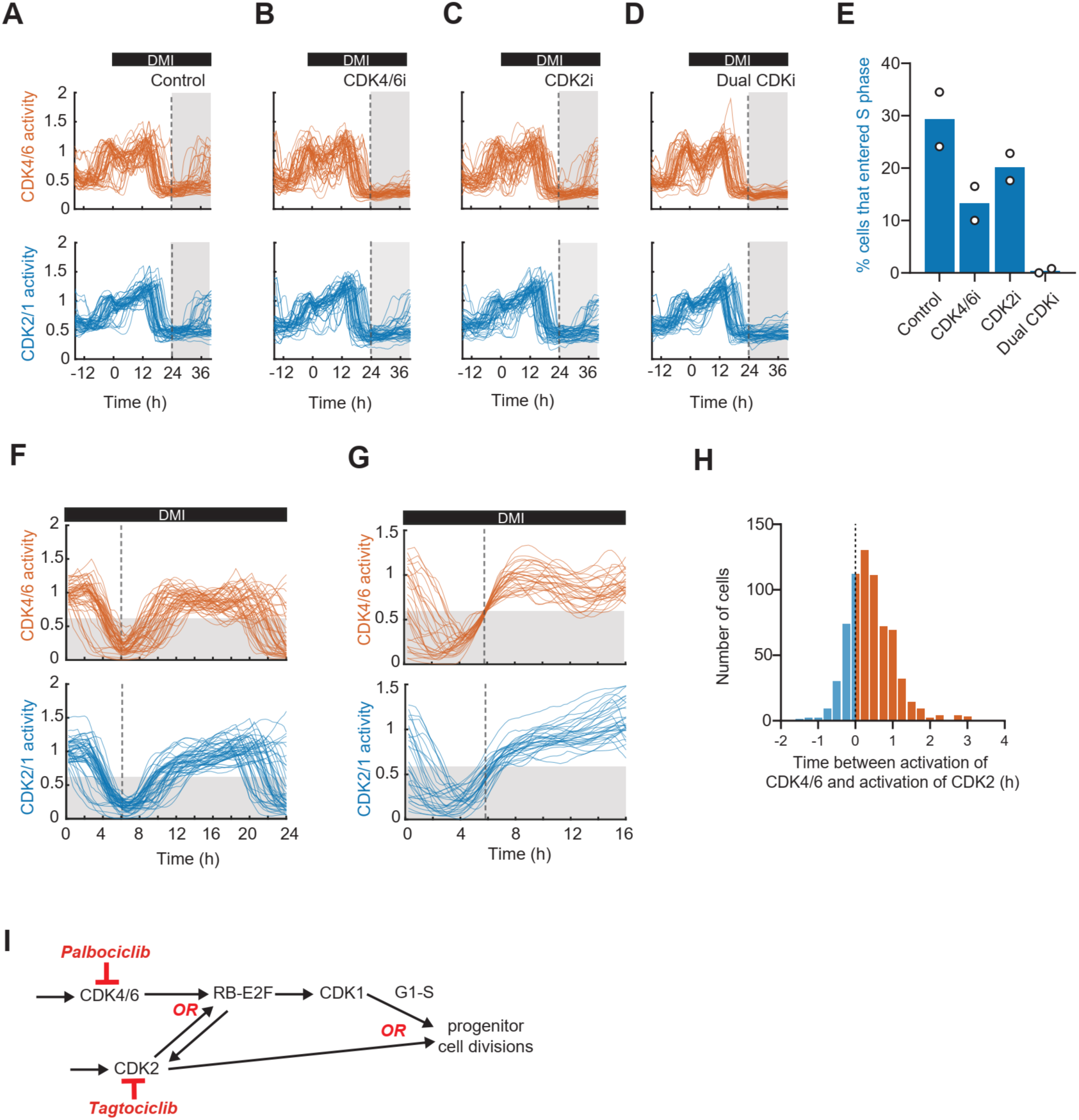
Progenitor cell divisions during adipogenesis are redundantly controlled by CDK4/6 and CDK2 activities. (A-D) DMSO (control), CDK4/6 inhibitor (CDK4i, Palbociclib, 1uM), CDK2 inhibitor (CDK2i, Tagtociclib, 1uM), or both CDK4/6 and CDK2 inhibitors (Dual CDKi, 1uM of each) were added into the culture media 24h after cells were induced to differentiate with DMI (dashed vertical line). Plots show fifty single-cell time courses from cells that had undergone at least one mitosis event (CDK2 activity >1) 12h prior to and were in G1-phase when drugs were added. (E) Percent of cells in Figures 2A-E in which CDK2 (blue) activities cross a value 0.8 within 16h of drug addition, representing cells that would transition from G1 to S-phase. Datapoints represent independent wells from the same experiment with over 200 single cells analyzed per well. (F) Single-cell time courses of CDK4/6 (top) and CDK2 (bottom) activities aligned by when CDK4 activity reaches its minimum after mitosis (vertical dashed line). Baseline value was subtracted from both CDK4 and CDK2 activity values to compare their activation kinetics. (G) Single-cell time courses of CDK4/6 (top) and CDK2 (bottom) activities aligned by when CDK4 activity increases to a value of 0.6 after the drop at mitosis. Baseline value was subtracted from both CDK4 and CDK2 activity values to compare their activation kinetics. After baseline correction, CDK activity of 0.6 represents the transition from G1 to S-phase. Figures 2F-G are representative of three biological replicates. (H) Distribution of difference in time for CDK4/6 activity versus CDK2 activity to reach a value of 0.6 in cells after mitosis in the presence of DMI. Blue bars represent cells where CDK2 activity rises before CDK4/6 activity to a value of 0.6 whereas orange bars represents where CDK4/6 activity rises earlier than CDK2 activity. Figures 2F-2G show data from the same experiment. (I) Progenitor cell divisions are driven by alternate routes through redundant CDK4/6 and CDK2 activation.

When analyzing the relative kinetics of CDK activity during G1 phase, we found that CDK4/6 activity increases slightly before CDK2 activity in most differentiating cells. We demonstrated this delayed activation of CDK2 using two distinct methods: aligning single-cell time courses to the minimum CDK4/6 activity after mitosis (Figure 2F) or to the timepoint at which CDK4/6 activity reaches a value of 0.6, representing the transition from G1 to S-phase (Figure 2G). Additionally, histogram analysis confirmed that most cells activate CDK2 only after CDK4/6 is activated (Figure 2H). Taken together, these results support that progenitor cells first activate CDK4/6, which then boosts CDK2 activity to drive the next cell cycle. Or if CDK2 is inhibited, CDK4/6 boosts CDK1 activity.

Initially, it was puzzling how cells lacking either CDK2 or CDK4/6 activity could enter S phase, as both kinase activities are often considered necessary for triggering cell cycle^20^. However, gene knockout studies of cyclins and CDK’s in mice^21–23^ support that CDK2 and CDK4/6 can, at least in some cells, on their own phosphorylate RB and activate the cell cycle transcription factor E2F (Figure 2i). Our data shows that both CDK4/6 and CDK2 can be independently activated in the progenitor cells when either is inactive. Furthermore, each can on its own activate E2F in G1 phase since many cells still enter the cell cycle when either CDK was inhibited (Figure 1J-K, 2B-C). Progenitors lacking CDK2 activity can still enter S phase data (Figure 1K and 2C), supporting that CDK4/6 activated E2F can activate CDK1, instead of CDK2, to initiate S phase (as diagrammed in Figure 2I), but only after an apparent delay. We thus conclude that there are two critical cell cycle redundancies in progenitor cells: progenitors can activate E2F in G1 phase by activating either CDK4/6 or CDK2, and can then enter S phase by either activating CDK2 or CDK1.

### PPARG induction requires that CDK4/6 and CDK2 are inactivated in G1 phase

We next used our triple-reporter cells to understand the relationship between CDK4/6 and CDK2 activities and the levels of PPARG during terminal differentiation. Figure 3A shows time courses of PPARG levels and CDK4/6 and CDK2 activities in cells that went on to differentiate in response to an adipogenic stimulus applied for the first 72 hours. We found that CDK4/6 and CDK2 were inactivated at approximately 18-36 hours, about the same time as PPARG reached the differentiation commitment threshold (Figure 3A, dashed line). We note that the differentiation protocol includes withdrawal of DMI after 3 days and replenishment of fresh insulin-containing media, which resulted in a small increase of CDK4/6 and CDK2 activities at 72 hours that did not trigger cell cycle entry (Figure 3A). Such a reversible partial CDK activation without cells starting the cell cycle has previously been characterized in other cells in response to weak mitogenic stimuli^24^.

**Figure 3.**
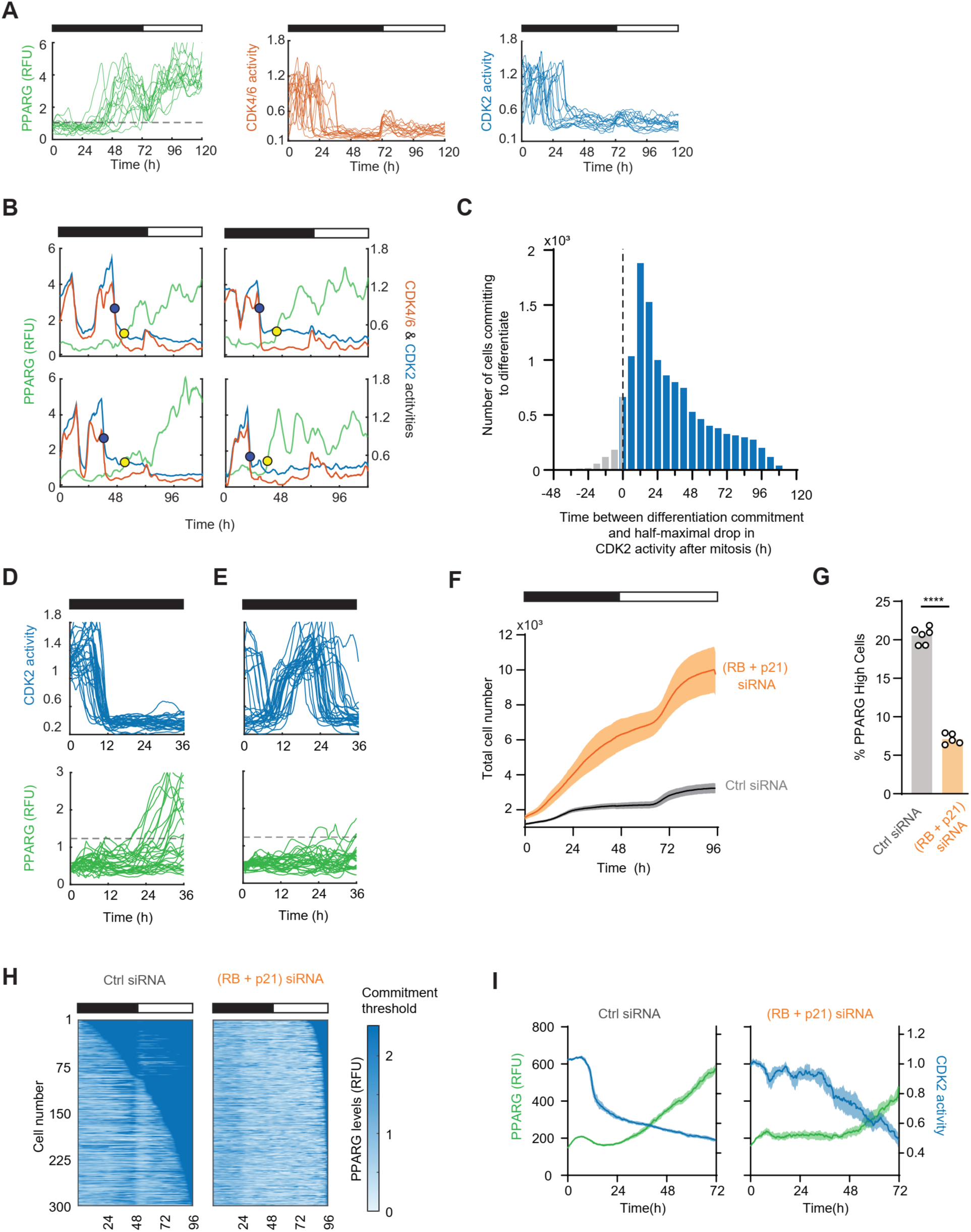
PPARG induction requires inactivation of CDK4/6 and CDK2 in G1 phase. (A) Triple reporter cells expressing endogenous mCitrine-PPARG, CDK4/6 and CDK2 reporters were induced to differentiate with a DMI stimulus for 72h (black horizontal bar), imaged and tracked over 5 days. Single-cell time courses of mCitrine-PPARG (left), CDK4/6 (middle) and CDK2 (right) activities from cells that increase PPARG levels beyond the commitment threshold. Representative of >10,000 cells tracked in the experiment. (B) Representative single cell time courses of mCitrine-PPARG (green) and CDK4/6 (Orange), and CDK2 (blue) reporters from differentiating cells. The yellow circle marks the time when cells reach the PPARG threshold (differentiation commitment timepoint). The blue circle marks the end of mitosis, as defined by when the CDK2 activity has dropped by more than half its peak level before mitosis. CDK4/6 activity is also seen to drop closely with CDK2 activity. (C) Histogram of the time between the end of mitosis (when CDK2 is inactivated) and the differentiation commitment time for all cells that differentiated. (D-E) 30 single cell time courses of CDK2 activity (top) and mCitrine-PPARG (bottom) from cells that have suppressed (D) or those that continue to increase CDK activity (E) between 12 and 36h after DMI addition. Dashed line indicates the PPARG commitment threshold. Data for Figure 3A-3E are from the same experiment and representative of three biological replicates. (F) Time course of total number of cells that were tracked in control and RB/p21 double knockdown conditions. Plot shows mean (line) ± SEM (shaded regions) from 5-6 replicate wells. (G) mCitrine-PPARG cells were treated with control or RB and p21 siRNA and imaged live during a standard 96h DMI induced differentiation protocol. Percent of cells with PPARG levels higher than the different commitment threshold at 96h was calculated (see Figure1B and Methods) for control and RB/p21 knockdown conditions. Bars show the mean and individual datapoints from 5-6 replicate wells in the experiment. Representative of three biological replicates. Unpaired two-tailed t-test used to compare means of control and RB + p21 knockdown conditions, ****P<0.0001. (H) Heatmap plots from three hundred control and RB + p21 knockdown cells, showing the times at which cells increase PPARG and commit to differentiation (seen as the deep blue regions on the plots). Representative of three biological replicates. (I) Time course of mCitrine-PPARG levels and CDK2 activity from triple reporter cells that start with high CDK2 activity (greater than 1) and eventually increase PPARG levels above the commitment threshold from control and RB+p21 knockdown conditions. Lines show median and the shaded regions represent 95% confidence intervals around the median. (Ctrl siRNA – 680 cells, RB + p21 siRNA – 124 cells). Representative of three biological replicates.

Figure 3B shows representative time courses from individual cells in which the drop in CDK activity after mitosis is marked with a blue dot and the PPARG threshold is marked with a yellow dot. These time courses, together with a histogram analysis in Figure 3C, support that the inactivation of CDK4/6 and CDK2 precedes the increase in PPARG. Nearly all cells increase PPARG after CDK2 inactivation (Figure 3D), but markedly, the PPARG increase happens with variable delays that can last for days. In cells in which the CDKs were inactivated early after adipogenic stimulation (<12 hours, Figure 3D), PPARG generally increased early, while PPARG stayed low in cells that inactivated CDKs late (Figure 3E), further suggesting that inactivation of the CDKs is required for PPARG levels to increase.

To determine whether CDK4/6 and CDK2 inactivation is required before cells can increase PPARG and initiate differentiation, we used siRNA to deplete p21 and RB, two critical suppressors of CDK4/6 and CDK2^7^, and thereby force cells to keep proliferating despite the presence of the differentiation stimulus. Indeed, increasing CDK4/6 and CDK2 activities during adipogenesis by depletion of p21 and RB resulted in increased proliferation, as evidenced by the strong increase in total cell numbers (Figure 3F), while at the same time greatly reduced the fraction of cells that reached the PPARG threshold (Figure 3G). Moreover, the few cells that eventually committed to differentiate only did so after a long delay (Figure 3H), corresponding to the time when CDK2 activity started to become inhibited (Fig 3I).

We conclude that differentiating progenitor cells decide in each G1 phase whether to enter an additional cell cycle by activating CDK4/6 and/or CDK2. However, they only undergo a limited number of divisions (from 1 to about 4, shown earlier in Figure 1C) before invariably exiting the cell cycle, which requires that they permanently inactivate both CDK4/6 and CDK2 in G1 phase. Furthermore, active CDK4/6 and CDK2 suppress the PPARG increase and CDK4/6 and CDK2 must both be inactivated before progenitors can increase PPARG and commit to differentiate.

### PPARG-induced p21 and p18 enforce the cyclin D1 and p27-regulated CDK4/6 and CDK2 inactivation

Our results in Figure 3A-D demonstrated that CDK4/6 and CDK2 inactivation precedes differentiation commitment and that PPARG only increases after cells stay for prolonged periods in G1 without CDK4/6 and CDK2 activities. Previous studies showed that PPARG directly drives p21 expression transcriptionally^5,25^, explaining how cells can permanently exit the cell cycle after PPARG is allowed to increase and reach the threshold. However, what regulators inactivate CDK4/6 and CDK2 early in the terminal differentiation process when PPARG levels are low was not clear. We therefore examined how the levels of cyclin D1 (CCND1), p21, p27 and p18, the four main regulators of CDK4/6 and CDK2 activity in G1, change during the time course of adipogenesis, comparing their levels in single cells that are specifically in G1-phase and have low or high PPARG.

When we measured the nuclear levels of cyclin D1 and p27 in individual G1 cells as a function of time, we found a gradual reduction of cyclin D1 and increase in p27 in both PPARG-high and PPARG-low cells, arguing that the expression of these two regulators is not dependent on PPARG levels (Figure 4A, B, see also Supplementary Figure 2). Markedly, cyclin D1 levels decreased and p27 levels increased already at 24 hours, before progenitor cells typically increase PPARG levels with the DMI stimulus. The decrease in cyclin D1 can explain the lower CDK4/6 activity but the increase in p27 is surprising since p27 has been proposed in some studies to activate CDK4/6 activity^26^. However, one recent study suggest that localization is critical and the nuclear level of p27 (which we measure here) is primarily inhibiting both cyclin D-CDK4/6 and cyclin E-CDK2 activity^27^. Thus, the combined decrease in nuclear cyclin D1 and increase in p27 can explain the inhibition of CDK4/6 and CDK2 activity and lengthening of the G1 phase before PPARG increases.

**Figure 4.**
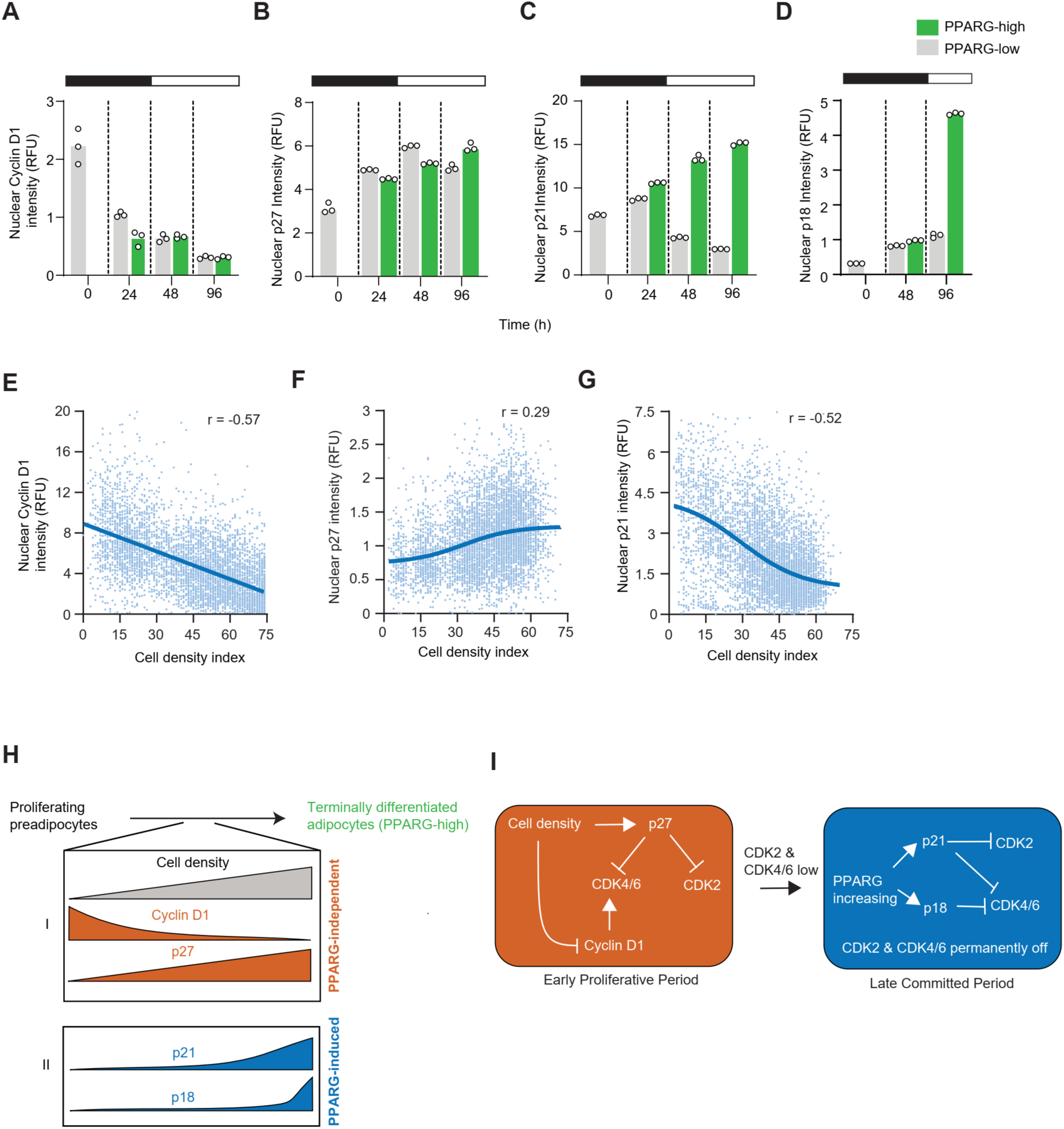
PPARG-induced p21 and p18 solidify cyclin D1 and p27-regulated CDK4/6 and CDK2 suppression. (A-D) OP9 cells expressing endogenous mCitrine-PPARG were plated across multiple 96 well plates in parallel, stimulated with DMI to induce differentiation and a plate was fixed at each of the indicated timepoints for single-cell immunofluorescence analysis. Immunofluorescence measurements of nuclear cyclin D1 (A), nuclear p27 (B), nuclear p21 (C) and nuclear p18 (D) levels from cells that have PPARG levels higher (green) or lower (grey) than the differentiation commitment threshold at every timepoint. Plots show overall means and the individual means of 3 replicate wells in the experiment wherein at least 1000 cells were analyzed per well. Representative of two biological replicates. (E-G) Scatter plot of nuclear cyclin D1 (E), p27 (F) and p21 (G) levels versus local cell density from single unstimulated OP9 cells in culture, measured 24h after they were plated at different densities. Line on each plot shows a least squares regression fit to the datapoints. Pearson’s correlation co-efficient (r) for the fitted line is indicated on each plot. Representative of two biological replicates. (H) As proliferating preadipocytes undergo differentiation, the increasing cell density causes cyclin D1 and p21 levels to decrease and p27 levels independently of PPARG expression thereby suppressing CDK activation across all cells. Additionally, in cells that increase PPARG levels in the presence of an adipogenic stimulus, PPARG drives the expression of p21 and p18 to maintain permanent cell cycle arrest. (I) Cell density causes a reduction in cyclin D1 and an increase in p27 levels to suppress CDK4/6 and CDK2 activity as progenitors proliferate. Increasing PPARG expression in the differentiation committed state permanently shuts off CDK4/6 and CDK2 activities through expression of p21 and p18.

In contrast to the sustained increase in p27 at 24 hours in all cells, p21 levels dropped in progenitors that did not increase PPARG, suggesting that p21 does not contribute to the initial inactivation of CDK4/6 and CDK2 activity (Figure 4C). However, p21 greatly increased in cells with high PPARG (Figure 4C), consistent with p21 being induced by PPARG^5^. This late induction suggests that p21 only inactivates CDK4/6 and CDK2 after cells have committed to differentiate. Moreover, the CDK4/6 inhibitor p18, which also has been implicated in many differentiation processes, was greatly increased by PPARG but after an even longer delay (Figure 4D) and is therefore contributing later to the permanent cell cycle arrest.

Studies in other cell types have shown that increasing cell density reduces cyclin D1 levels and increases p27 levels^27^,and we evaluated if this also occurs during differentiation. Such a mechanism seemed plausible since progenitor cell density has been shown to be a critical parameter to enhance adipogenesis^5,28^. Indeed, increasing the local progenitor cell density reduced nuclear cyclin D1 and increased p27 levels (Figure 4E, F), explaining how increased progenitor cell density during differentiation can inactivate CDK4/6 and CDK2. In contrast, p21 levels were decreased rather than increased at higher progenitor density (Figure 4G), further arguing that it is the reduction of cyclin D1 and increased p27 rather than the change in p21 that is mediating the initial inhibition of CDK4/6 and CDK2 activity during differentiation.

We conclude that adipogenic stimulation gradually reduces the nuclear cyclin D1 level and increases the nuclear p27 levels through a cell division-mediated increase in progenitor density, which inactivates both CDK4/6 and CDK2 to lengthen the G1 phase independent of PPARG (Figure 4H). The lengthened G1 then gives PPARG sufficient time to increase to the commitment point and trigger the second step, the induction of p21 and later p18, which permanently suppress CDK4/6 and CDK2 activity in the terminal differentiation state (Figure 4H). Mechanistically, p21 and p27 can inhibit both cyclin E-CDK2 and cyclin D-CDK4/6 complexes while cyclin D1 activates and p18 inhibits CDK4/6 alone (Figure 4i).

### Delayed cell density and DMI-mediated ERK inactivation is also required for PPARG to increase

Since CDK2 and CDK4/6 inactivation is necessary for differentiation commitment, we next tested whether their inactivation is also sufficient to induce differentiation. To do so, we determined whether premature inhibition of CDK4/6 and CDK2 promotes differentiation. Control experiments confirmed that combined CDK4/6 and CDK2 inhibition lowered the percent of cells that proliferated, as measured by the presence of APC/C reporter fluorescence (Figure 5A, Supplementary Figure 4A) and by the presence of phosphorylated Rb (Figure 5B, Supplementary Figure 4B). Unexpectedly, when we inhibited CDK4/6 and CDK2 (dual CDKi) after 24 hours, there was only a small increase in the percentage of differentiated cells (Figure 5C), indicating that CDK4/6 and CDK2 inactivation is not the main factor restricting differentiation commitment.

**Figure 5.**
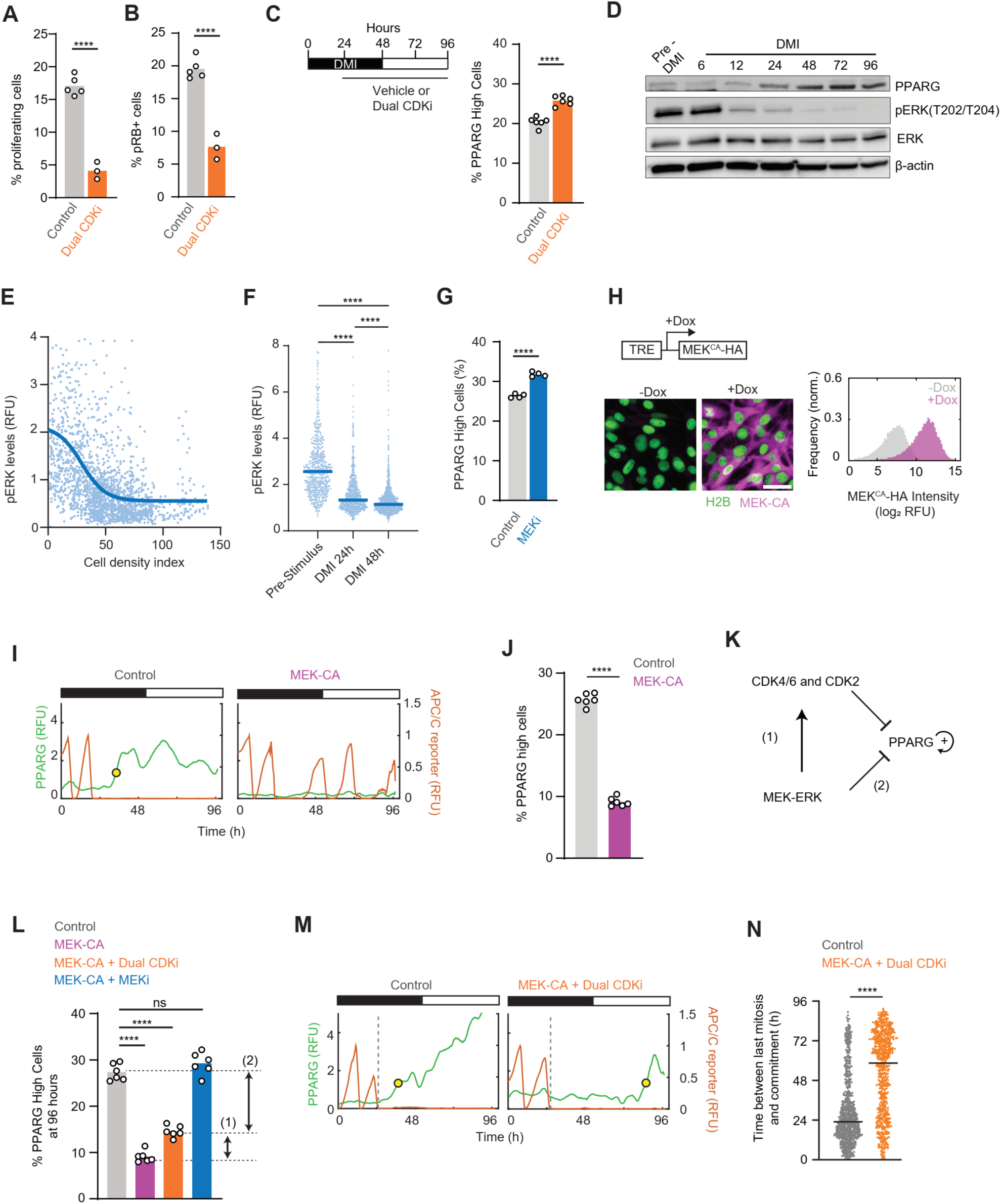
PPARG induction requires CDK4/6 and CDK2, as well as ERK, inactivation. (A-B) Proliferating OP9 preadipocytes expressing APC/C-reporter cell cycle reporter were treated with a combination of Palbociclib and CDK2i (1 μM each) for 24h, fixed and immunostained for phosphorylated RB protein levels. Plots show mean and the individual datapoints from 3-5 replicate wells in the experiment. Representative of two biological replicates. (Also see Supplementary Figure 4) (A) Percent of proliferating cells (i.e. cells in S/G2/M phases) defined by the number of cells positive for nuclear APC/C reporter fluorescence. (B) Immunofluorescence staining was used to measure the percent of cells positive for phosphorylated RB (pRB). (C) mCitrine-PPARG cells were imaged live during a standard 96h DMI induced differentiation protocol and CDK inhibitors (or vehicle) were added and kept in starting 24h after the DMI stimulus (experimental scheme, left panel). Dual CDKi refers to co-inhibition with Palbociclib and Tagtociclib, 1uM each. Percent of cells with PPARG levels higher than the different commitment threshold at 96h was calculated (see Figure1B and Methods) for control and Dual CDKi-treated conditions (right panel). Plots show the mean and individual datapoints points from 6 replicate wells in the experiment. Representative of three biological replicates. (A-C) Unpaired two-tailed t-test was used to compare sample means. ****P<0.0001. (D) Western blot analysis for phospho-ERK (pERK) and total ERK and PPARG levels during adipogenesis. Beta-actin was used a loading control. Data is representative of two biological replicates. (E) Scatter plot of pERK levels (nuclear + cytosolic) levels versus local cell density from single unstimulated OP9 cells in culture, measured 24h after they were plated at different densities. Line on each plot shows a least squares regression fit to the datapoints. Representative of two biological replicates. (F) Distribution of pERK from single OP9 cells at similar densities (cell density index of 30-45) before, at 24h, and 48h of DMI stimulation. One-way ANOVA with Tukey’s multiple comparisons was used to compare the difference between conditions. ****P<0.0001. More than 500 cells analyzed per condition. (G) mCitrine-PPARG cells were imaged live during a standard 96h DMI induced differentiation protocol and MEK inhibitor (PD0325901 – 100nM) or vehicle was added and kept in starting 24h after the DMI stimulus. Percent of cells with PPARG levels higher than the different commitment threshold at 96h was calculated (see Figure1B and Methods) was calculated for control and MEKi-treated conditions. Plots show the mean and individual datapoints points from 4 replicate wells in the experiment. Representative of three biological replicates. Unpaired two-tailed t-test was used to compare differences between sample means. ****P<0.0001. (H) Scheme for the doxycycline-inducible MEKCA construct (top left). Immunofluorescence images (bottom left) and distribution of HA-tagged MEKCA expression (right) in control and doxycycline-induced cells after 24h of induction. Scale bar - 25 μm. (I-K) mCitrine-PPARG cells were treated with doxycycline (2μg/mL) to induce the expression of MEKCA (or vehicle) during a standard 96h DMI induced differentiation protocol DMI stimulation and imaged over 96h. (I) Representative single cell time courses of PPARG and APC/C cell cycle reporter from control and MEK-CA overexpressing cells. Yellow circle shows the differentiation commitment point. (J) Percent of cells with PPARG levels higher than the different commitment threshold at 96h was calculated (see Figure1B and Methods) was calculated for control and MEKCA overexpression conditions. Plot shows means and individual datapoints from 6 replicates wells in the experiment. Representative of two biological replicates. Unpaired two-tailed t-test was used to compare differences between sample means. ****P<0.0001. (K) MEK-ERK signaling may suppress the rise in PPARG both through promoting CDK activity and by directly phosphorylating and suppressing PPARG transcriptional activity and prevent any positive feedback on its expression levels. (L) Besides doxycycline for MEKCA expression, cells were also treated with vehicle, Dual CDKi (Palbociclib - 4 μM + CDK2i - 1μM) or MEKi (PD0325901 - 100nM) or Dual CDKi + MEKi. Inhibitors (or vehicle) were added and kept in starting 24h after the DMI stimulus. Percent of cells with PPARG levels higher than the different commitment threshold at 96h was calculated (see Figure1B and Methods) for control and inhibitor(s)-treated conditions. Plot shows means and individual datapoints from 6 replicates wells in the experiment. Representative of two biological replicates. One-way ANOVA with Tukey’s multiple comparisons test was used to compare differences between conditions. ****P<0.0001, ns – not significant. (M) Representative single cell time courses of PPARG and APC/C cell cycle reporter from control and MEK-CA cells treated with Dual CDKi (Palbociclib - 4 μM + CDK2i - 1μM) as described for Figure 5L, showing variability in the timing of commitment to differentiation (yellow circle) after completion of mitosis (dashed line). (N) Distribution of the time taken after mitosis to undergo differentiation commitment from single differentiating cells for experiment represented in Figure 5M. Horizontal line shows the median of each distribution. Mann-Whitney test used to compare medians of control and MEK-CA + CDKi conditions, control - 770 cells, MEK-CA + CDKi - 655 cells. Representative of two biological replicates. ****P<0.0001. Examples traces in Figure 5I and 5L from the same experimental dataset.

We hypothesized that this unknown suppressive factor might be MEK/ERK kinase activity since ERK can phosphorylate PPARG to suppress the transcriptional activity of PPARG^29^. However, there have been conflicting reports whether ERK promotes or inhibits adipogenesis^29–31^, leaving its role unresolved. We considered that these contradictory results might be explained by the timing when MEK/ERK is active during differentiation. We hypothesized that MEK/ERK signaling may be needed early in differentiation to promote proliferation but must then be inactivated to allow for PPARG to increase the PPARG levels. This dual role seemed plausible, since MEK/ERK activity also promotes proliferation by inducing cyclin D1 expression^32^. Indeed, ERK activity was progressively reduced during differentiation as evidenced by the reduction of phospho-ERK levels by western blot analysis (Figure 5D).

We identified two contributing factors that inactivate ERK during adipogenesis: First, the early period of differentiation is paralleled by increasing progenitor density due to the ongoing cell divisions. Analysis of ERK activity as a function of the local progenitor density showed that increasing the local density suppresses ERK activity (Figure 5E). An analogous cell density-mediated ERK inactivation has been characterized in other cell types^33^. Second, when we analyzed ERK phosphorylation as a function of time after adipogenic stimulation by selecting progenitors with the same local density, we found that ERK is further inhibited by the adipogenic stimulus independent of local progenitor density (Figure 5F). This inhibition of ERK can potentially be explained by the upregulation of the ERK phosphatase DUSP1 by glucocorticoids since a glucocorticoid is part of the differentiation stimulus^34^. Crucially, when we treated preadipocytes with MEK inhibitor (MEKi) 24 hours after DMI stimulus was added, a higher percent of cells increased their PPARG levels after MEK inhibition (Figure 5G), demonstrating that ERK inactivation promotes the increase in PPARG and differentiation commitment.

To directly determine whether ERK inactivation is necessary for the PPARG increase and commitment to differentiate, we generated a Doxycycline (Dox)-inducible CA-MEK construct to prolong ERK activity during adipogenesis^27^. We stably introduced this construct into OP9 preadipocyte cells expressing mCitrine(YFP)-PPARG (Figure 5H, see also Supplementary Figure 3). Control experiments showed that the CA-MEK induction prolonged cell proliferation (Figure 5I).

Markedly, addition of Dox to induce CA-MEK and persistently activate ERK strongly inhibited the PPARG increase (Figure 5I, J). Since CDK4/6 activation also inhibits differentiation, this raises the question whether ERK inhibits PPARG indirectly through activating CDK4/6 (Figure 5K, pathway 1) or directly through inactivating PPARG (Figure 5K, pathway 2). To test which of these mechanisms is critical, we inhibited CDK4/6 and CDK2 in CA-MEK expressing cells which only partially restored the percent of differentiating cells. However, additional MEK inhibition restored the fraction of differentiation cells seen in the control conditions (Figure 5L). In further support that ERK activity has a direct repressive role independent of CDK4/6 and CDK2, differentiation commitment in all MEK-CA cells with inhibited CDK4/6 and CDK2 was greatly delayed (Figure 5M, N).

We conclude that ERK must be inactivated to allow progenitors to increase PPARG levels and commit to differentiate. Inactivation of ERK is redundantly controlled by the adipogenic stimulus and increasing local progenitor density by the ongoing cell divisions in the proliferative period. ERK activity is inhibiting differentiation commitment both indirectly by activating CDK4/6 and directly by preventing PPARG levels to increase.

### Sequential inactivation of CDK4/6-CDK2 and ERK separates the reversible proliferative period from the commitment to differentiate

Combined inhibition of CDK2, CDK4/6 and MEK 24 hours after adipogenic stimulation had an additive effect compared to the inhibition of the two CDKs or MEK alone (Figure 6A), suggesting that different cells in the population may be relying on prolonged ERK or CDK activation to suppress PPARG and differentiation. To directly determine whether inactivation of the CDKs or inactivation of ERK is the rate-limiting step for increasing PPARG, we monitored the time course of the PPARG increase after adding MEK or CDK inhibitors 24 hours after the DMI stimulus. Markedly, MEK inhibited cells rapidly increased PPARG levels compared to control cells (Figure 6B), while inhibition of the CDKs only increased PPARG after a delay. Moreover, combined CDK and MEK inhibition increased PPARG levels much more rapidly than CDK inhibition alone, implying that inhibition of ERK activity is generally the rate limiting step acutely increasing PPARG (Figure 6C). This acceleration of the PPARG increase by MEK inhibition is more clearly shown when comparing the averaged signals in Figure 6D and E. The accelerated increase in PPARG by MEK inhibition can also be shown in individual cells in a heat map analysis by sorting 500 progenitors by the time when PPARG increases and comparing them under MEK and CDK inhibited conditions (Figure 6F and G). We conclude that ERK inactivation is an acute trigger increasing PPARG levels in cells that have already inactivated CDKs.

**Figure 6.**
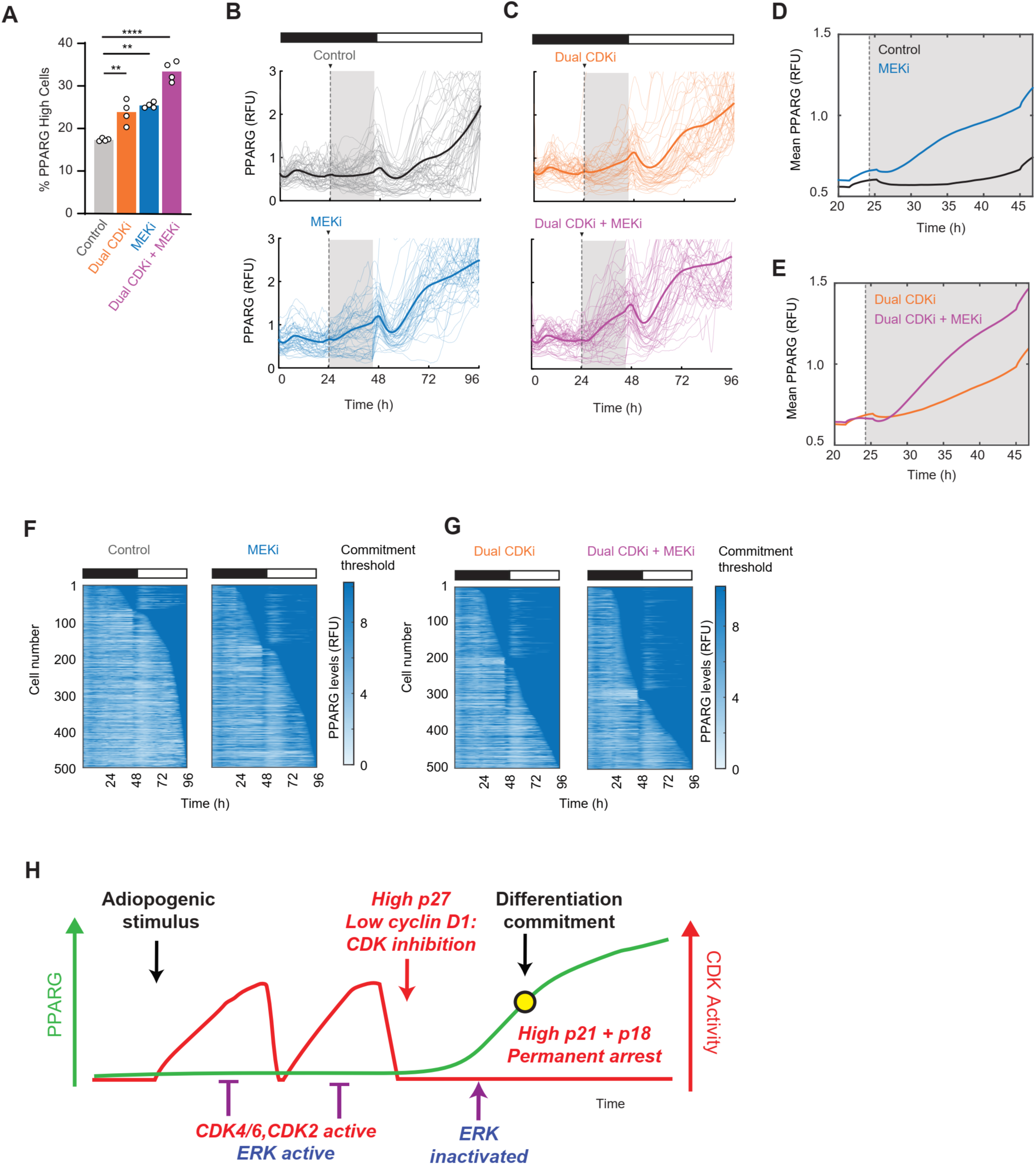
Sequential inactivation of CDK4/6-CDK2 and ERK separates a reversible proliferative period from commitment to differentiation. (A) mCitrine-PPARG cells were imaged live during a standard 96h DMI induced differentiation protocol and inhibitors, Dual CDKi (Palbociclib - 4 μM + CDK2i - 1μM), MEKi (PD0325901 - 100nM), Dual CDKi + MEKi or vehicle were added and kept in starting 24h after the DMI stimulus. Percent of cells with PPARG levels higher than the different commitment threshold at 96h was calculated (see Figure1B and Methods) was calculated for control and inhibitor(s)-treated conditions. Plot shows the means and individual datapoints from 4 replicate wells in the experiment. One-way ANOVA with Dunnett’s multiple comparisons test was used to compare differences between conditions. **P< 0.01, ****P<0.0001. (B-C) 50 single cell time courses of mCitrine-PPARG from cells that eventually commit to differentiate across the control and MEKi (B) and, Dual CDKi and Dual CDKi + MEKi (C) treatment conditions. The bold line shown the mean PPARG levels. Representative of >500 cells tracked per condition in the experiment. Shaded regions highlight the differences seen within 24h of drug addition. (D-E) Mean mCitrine PPARG levels for control and MEKi (D) and Dual CDKi and Dual CDKi + MEKi (E) conditions 4 hours prior to and for 24h (shaded regions in Figure 6B and 6C) after the drugs were added. (F-G) Heatmap plots from control and MEKi (F) and Dual CDKi and Dual CDKi + MEKi (G) treatment conditions, showing the times at which cells increase PPARG and commit to differentiation (seen as the deep blue regions on the plots). Data in Figures 6A-6G are from the same experiment and are representative of three biological replicates. (H) Progenitor cells exposed to differentiation stimulus undergo divisions as long as CDK4/6 and CDK2 activity increase during G1 phase. Active MEK-ERK signaling in G1-phase drives cyclin D1 expression and CDK activity. ERK also phosphorylates and suppresses the transcriptional activity of any PPARG expressed in cells, preventing any downstream positive feedback that increase PPARG expression. As progenitors proliferate, increasing cell density and reduction in ERK activity cause a decrease in cyclin D levels and an increase in p27 levels to suppress CDK4/6 and CDK2 activation in the G1 phase. This induces a reversible state of cell cycle arrest until further ERK inactivation increases PPARG transcriptional activity, which both induces differentiation commitment and increases p21 and p18 levels to enforce permanent cell cycle arrest. Thus, inactivation of both CDK and ERK activities in G1-phase is required for cells to commit to differentiate and establish a post-mitotic state. Extending activation of either CDK or ERK signaling delays or prevents differentiation.

## DISCUSSION

Our study investigated how cell cycle exit is coordinated with the commitment to terminal differentiation. We employed live single-cell reporter analysis to directly elucidate the relationship between PPARG, the primary transcriptional regulator of adipogenesis commitment, and the two critical G1-phase kinases, CDK4/6 and CDK2, which control cell cycle entry and exit. We demonstrated that redundant activation of CDK4/6 and CDK2 in G1 phase drives the proliferative phase of adipogenesis, allowing for a variable but limited number of cell divisions (Figure 6H). Following each mitosis, CDK4/6 and CDK2 activities decrease, providing cells the opportunity to decide between proceeding to the next cell cycle or exiting. Only cells that maintain CDK4/6 and CDK2 inactivity for sufficient durations can increase PPARG to the threshold where progenitors commit to terminally differentiate.

We identified two sequential steps leading to the permanent inactivation of CDK4/6 and CDK2 during adipogenesis. Initially, increased local progenitor density, resulting from ongoing proliferation, transiently inhibits CDK4/6 and CDK2 by reducing cyclin D1 and increasing p27 levels. Subsequently, PPARG-mediated induction of p21 and p18 permanently inhibits CDK4/6 and CDK2 after cells commit to differentiation. Surprisingly, inactivation of CDK4/6 and CDK2 alone was insufficient to induce the PPARG increase and commitment to differentiation. Cells could only increase PPARG and commit to differentiation if they also inactivated ERK, which we identified as the rate-limiting step initiating differentiation commitment.

Our findings expand upon previous research demonstrating that progenitor cells permanently exit the cell cycle during terminal differentiation and that proliferation and differentiation mutually suppress each other^1–3^. However, due to significant variability in the timing of individual cell cycle exit and differentiation initiation^5^, a mechanistic understanding of how these processes are coordinated has remained elusive without live-cell analysis. Our adipogenesis cell model, incorporating endogenously tagged PPARG and mosaic time-course analysis, enabled us to precisely determine the differentiation commitment point while simultaneously monitoring the activity of both CDK4/6 and CDK2 relative to differentiation commitment during the 4-to-5-day differentiation process.

Previous studies have reported conflicting roles of ERK activity in promoting or suppressing adipogenesis. A study by Farmer’s group concluded that ERK activity is required for adipogenesis, while Spiegelman’s group demonstrated that ERK activity directly phosphorylates PPARG and inhibits its transcriptional activity^29,30^. It was also plausible that MEK-ERK activity might indirectly suppress PPARG by activating CDK4/6, since MEK-ERK activity has been shown to induce Cyclin D expression^32^.

Our study supports that ERK may be both promoting and suppressing adipogenesis, but just at different times in the process: First, ERK activity, by inducing Cyclin D expression^32^, activates CDK4/6 and drive cell cycle progression early in differentiation. Second, we show that ERK must be inhibited late in differentiation, not only to inhibit CDK4/6 but also to allow PPARG to increase independently of CDK4/6 and CDK2 (summarized in Figure 6H). Furthermore, we show that the delayed inactivation of ERK arises both from the increase in local progenitor density due to the cell division early in adipogenesis and from the adipogenic stimulation itself. Thus, ERK activity plays a critical role in promoting proliferation early in adipogenesis, but must subsequently be inhibited to both inactivate CDK4/6 and allow PPARG to increase.

A previous study showed that knocking out the cell cycle inhibitors p21 and p27 in mice greatly increased the number of differentiated cells (hyperplasia) and caused obesity^9^. Our study suggests that this adipose tissue hyperplasia results from a prolonged proliferative period early in the terminal differentiation process. This prolonged proliferative period, driven by the absence of p21 and p27, allows for the generation of more adipocytes per progenitor cell.

The regulatory mechanisms governing adipogenesis may also explain how proliferation and differentiation are interconnected in other cellular systems, such as neurons and muscle. In these systems, progenitor cells initially undergo expansion through repeated cell divisions before daughter cells exit the cell cycle and terminally differentiate^2,10^. Similar to PPARG inducing p21 in adipocytes, the main transcriptional drivers of differentiation, NeuroD and myoD, also induce p21 in neurons and muscle, respectively. Given the general role of ERK, CDK4/6, and CDK2 in promoting proliferation, it is plausible that external differentiation stimuli and local cell density generally first activate these proteins to control the length of the proliferative period. This proliferative period may not only facilitate epigenetic changes needed for differentiation, but also regulate the extent of regenerative responses by controlling the number of differentiated cells produced from each progenitor cell

Together, these results suggest the following framework for terminal cell differentiation: stimulated progenitors start with a variable proliferative period initiated by ERK, CDK4/6 and CDK2 activation, which must subsequently be terminated by reduced nuclear cyclin D1 and elevated p27 levels, leading to the inactivation of CDK4/6 and CDK2. In addition to this transient cell cycle arrest in G1, ERK activity must subsequently be inhibited to trigger differentiation commitment, which ultimately locks cells in permanent cell cycle arrest through the expression of p21, alongside other CDK inhibitors such as p18.

## METHODS

### Cell lines and constructs

The CDK1/2 (pLV-EF1a-DHB-mTurquoise) and CDK4/6 (pLV-EF1a-mCherry-KTR-RbCterm) reporter constructs were generated and characterized in Spencer et.al. 2013 and Yang et.al, 2020, respectively^15,16^. MEK1-CA construct was generated as described in Fan et.al, 2020^27^. Third-generation lentiviral packaging system was used to generate lentiviruses for all the reporter constructs as well as the fluorescently tagged nuclear reporter H2B constructs (H2B-mTurquoise and H2B-iRFP670) and used to infect and stably express these reporters in the OP9 preadipocyte cell line with endogenously tagged citrine-PPARG2. The dual-reporter cell line with endogenously tagged citrine-PPARG2 and stably expression of mCherry-Geminin-degron reporter was generated and characterized in Zhao et. al., 2020^5^. For all cell lines, either FACS was used to select and enrich for reporter positive cells or cells were subjected to antibiotic selection post lentiviral infection.

### Cell culture and differentiation

OP9 cell lines were cultured as previously published^35,36^. Briefly, the cells were cultured in phenol red containing MEM-α media (Thermo Fisher Scientific) supplemented with 20% FBS, 100 units/mL of Penicillin, 100 µg/mL Streptomycin and 292 µg/mL L-glutamine. Cells were induced to differentiate in the same basal media but supplemented with 10% FBS. In most experiments, a commonly-used DMI protocol was used induce adipogenesis, in which an adipogenic cocktail (DMI) consisting of dexamethasone (1 µM, Sigma-Aldrich), IBMX (125 µM, Sigma-Aldrich), and insulin (1.75 nM, Sigma-Aldrich) was added to the cell culture media for 48 hours, then aspirated away and replaced with fresh media containing 1.75 nM insulin for another 48 hours. Whenever experiments were performed with the triple reporter cell line expressing endogenous mCitrine-PPARG, CDK4/6 and CDK2 reporters, DMI containing media was kept on for 72 hours instead of the usual 48 hours. This extended stimulation time was used required to obtain a larger fraction of cells that go on to increase their PPARG levels beyond the differentiation threshold. For all live-cell microscopy experiments, cells were differentiated in the same media as described above but without phenol red.

### Inhibitors

The final concentrations for the small molecule chemical inhibitors used in this study were - CDK4/6 inhibitor Palbociclib (PD-0332991) – 1 or 4 µM as indicated. CDK2 inhibitor Tagtociclib (PF-07104091) - 1 µM, MEK inhibitor Mirdametinib (PD-0325901) - 100 nM. 4 µM Palbociclib was used in experiments where we drove the expression of constitutively active MEK and therefore expected basal CDK4 activity to be higher. For drug additions during adipogenesis, 40 μl of serum-free MEM-alpha media containing 5X concentration of the inhibitors was added to individual wells containing cells growing in 160 μl media, 24 hours after the DMI stimulus. Equal volumes of DMSO were used as vehicle control. Thereafter, media containing insulin and inhibitors, or vehicle was used to replace DMI containing media for the last 2 days of the differentiation protocol. All inhibitors were purchased from Selleck Chemicals.

### Fluorescent imaging

Cells were plated in full growth medium at a density of 20,000 cells/cm^2^ (for experiments at low density) 24h prior to imaging on to either CellVis (P96-1.5H-N) or Ibidi μ-Plates (catalog no. 89626). OP9 cells have a fibroblast-like morphology and show significant migratory behavior in culture, often transiently moving very close to or over neighboring cells. Therefore, for live-cell imaging experiments, to overcome the challenge of reliably identifying and tracking the same cells over several hours to days as they reach very high cell densities during the differentiation protocol, the cell suspension used for plating included a mixture of cells that either stably expressed fluorescently tagged H2B or not such we had a mosaic of labeled and unlabeled cells in every field of view being imaged. Ratio of tagged to untagged cells used was 1:1 to 3:1 (more labelled cells) whenever we expected perturbations to reduce proliferation of cells and 1:3 (more unlabeled cells) when the perturbations in the experiment was expected to increase proliferation. Before image acquisition, the full growth medium was aspirated and replaced with fresh MEMα without phenol red (R&D Biosystems, M34750) supplemented with 10% FBS containing the adipogenic stimulus wherever indicated. Live-cell imaging was performed on a Nikon ECLIPSE Ti2 inverted microscope using a 10× or 20x (for CDK reporter activity assays), Plan Apo 0.45-numerical aperture (NA) objective on a stage-top humidified 37 °C incubator maintained with 5% CO_2_. Images were acquired every 12-15 min for three or four fluorescent channels as the experiment required (CFP, YFP, RFP and iRFP). Total light exposure time was kept to less than 800 ms for each time point. Four to nine non-overlapping sites were imaged inside each well.

### siRNA-mediated gene silencing

siRNA targeting *p21, RB* and the AllStars Negative Control siRNA were purchased from QIAGEN. For siRNA knockdown in the live-cell imaging experiments, OP9 cells were transfected by reverse-transfection using Lipofectamine RNAiMax (Invitrogen). Briefly, our reverse-transfection protocol was: 19 µL of OptiMEM was mixed with 0.5 µL of a 10 µM siRNA stock solution, and 0.5 µL of Lipofectamine RNAiMax. Transfection mix was incubated at room temperature for 10 minutes and 80 µL of culture media was added which contained the desired number of cells per well. Then the entire (∼100µL) volume was plated into one well of a 96-well plate. The siRNA/RNAiMax mixture was left on the cells for 6-12 hours before being aspirated away and replaced with 100 µL fresh culture media. DMI was added 24h after siRNA transfection.

### Immunofluorescence (IF) staining

All cultured cells were fixed with 4% PFA in PBS for 15 min at room temperature, followed by three washes with PBS using an automated plate washer (Biotek). Cells were then pretreated with ice-cold methanol for 10 min, washed thrice with PBS and blocked for 1 hour in PBS containing 5% FBS and 0.3% Triton X-100. Cells were then incubated overnight with primary antibodies diluted PBS containing in 1% BSA and 0.3% Triton X-100 at 4°C. Primary antibodies used in this study were: mouse anti-PPARγ (Santa Cruz Biotech, sc-7273, 1:1,000), mouse anti-p21 (Santa Cruz Biotech, sc-6246, 1:100), rabbit anti-p21 (Abcam, ab-188224, 1:2000), mouse anti-p27 (Abcam, ab-193379, 1:500), rabbit anti-p27 (Abcam, ab-32034, 1:500), cyclin D1 (Abcam, ab-16663, 1:500), rabbit anti-p18 (Abcam, ab192239, 1:500), rabbit anti-HA (Cell Signaling Technology, 3274, 1:1000) and rabbit anti-phospho ERK1/2 (T202/Y204) (Cell Signaling Technology, 4370, 1:250). Following primary antibody incubation cells were washed with PBS (3X), incubated with Alexa fluor-conjugated anti-rabbit or anti-mouse secondary antibodies (1:1000, Thermo Fisher Scientific) in 1 PBS containing in 1% BSA and 0.3% Triton X-100 for 1.5 hours, followed by incubation with Hoechst (1:10,000 in PBS containing in 1% BSA and 0.3% Triton X-100) for 30 min at room temperature. Cells were then washed thrice with PBS using the automated plate washer prior to imaging.

### Image data processing and analysis

Data processing and analysis of fluorescent images was performed in MATLAB R2020a (MathWorks). Unless indicated otherwise, fluorescent images were processed, and intensity data was extracted by automated image segmentation, tracking and measurement using custom-written MACKtrack cell tracking software previously described in Kovary et al, 2018 and Zhao et al, 2020^5,37^. PPARG levels were quantified from fixed samples based on median fluorescence signal within the nuclei of individual cells. Cells were scored as ‘PPARG high’ if the marker expression level was above a cut-off determined from the bimodal distribution of PPARG expression at the end of the experiment.

For live imaging data acquired from OP9 cells, the CFP channel capturing H2B-mTurqoise or farred channel capturing H2B-iRFP670 fluorescence was used for nuclear segmentation and cell tracking. Obtained single-cell traces were filtered to remove incomplete or mis-tracked traces according to the following criteria: cells or their daughter cells that were absent at any point during the entire time lapse experiment, cells that had large increase or decrease in PPARG intensity normalized to the previous timepoint, cells where there are many and large fluctuations in H2B signal.

The percentage of PPARG high cells was calculated by counting cells in which the PPARG levels was above the threshold (calculated as described in the next section) at that time point as a fraction of the total number of cells being tracked at that time point.

To measure CDK (4/6 and 2) activity, the median fluorescence in the reporter channel was calculated for both the nuclear and cytosolic region around the nucleus. Nuclear fluorescence was calculated by directly using the H2B segmentation mask. For cytosolic fluorescence measurement, an annular region of interest around the nucleus was generated by isometrically expanding the nuclear segmentation mask by 4 pixels and deleting the nuclear mask from it and thereafter calculating the median reporter fluorescence within the annulus. Finally, the cellular CDK activity was calculated as the ratio of cytosolic to nuclear reporter fluorescence intensity for each cell at every timepoint.

Previous studies have shown that cells enter S-phase whenever the activity of CDK2 as measured by the respective reporters reaches a value between 0.8 and 1. We use the CDK2 activity value of 0.8 as a cut-off to identify cells that have entered S-phase (ref Sabrina’s 2012 Cell paper, Heewon’s 2020 Elife paper, Yumi’s 2024 Nature paper).

Local cell density measurements were performed by a method similar to that described in in Fan et.al, 2020^27^. Briefly, nuclear marker channel (DAPI or H2B) was used to segment the nuclei. Next, a centroid was estimated for every nucleus in the image. All nuclei (i.e. their centroids) that were less than ∼200 μm away from the edge of the image were excluded from further analysis. Finally, the number of neighboring centroids that were within ∼100 μm of every nucleus in all directions were counted as a measure of local cell density index.

### Estimating a differentiation commitment point (i.e. time at which cells increase PPARG above a threshold)

PPARG values at the end of a differentiation experiment typically exhibit a bimodal distribution when the differentiation stimulus is removed after 2 days. Under such conditions, a cut-off PPARG value (commitment threshold) was determined by scanning the entire PPARG time series data to estimate a PPARG value that identified the most number cells that would satisfy all of these three conditions: 1) PPARG levels at the start of the experiment was below threshold, 2) PPARG levels just prior to removal of adipogenic stimulus was above the threshold and 3) PPARG at the end of the experiment (96 or 120h) was above the threshold. Differentiation commitment time was then estimated for every cell that retained PPARG levels at or higher than this threshold at the end of the experiment by identifying when PPARG levels first cross this threshold during the experiment.

### Western Blotting

OP9 cells were plated in parallel on multiple 60 mm cell culture dishes in growth media at a density of approximately 20,000 cells/cm^2^ and stimulated with an adipogenic stimulus starting 48h after plating. At each time point indicated, cells on a single plate were washed twice with ice-cold PBS, scraped-off and lysed in ice-cold RIPA lysis buffer (Millipore Sigma, 20-188) containing a cocktail of protease and phosphatase inhibitors (Thermo Fisher Scientific, 78444). The lysates were incubated on ice for 20 min and spun down at 13000g for 10 min. The supernatant was collected, and total protein amounts for each sample were determined using 5 µL with a standard Bradford assay. 35-40 µg protein was used per sample for SDS-PAGE and western blotting. Blots were blocked in TBS containing 0.1% Tween-20 (TBST) and 10% dry milk and incubated overnight with primary antibodies diluted in TBST with 5% BSA. The following primary antibodies were used: rabbit anti-PPARG (Cell Signaling Technology, 2443, 1:1000), rabbit anti-phospho ERK1/2 (T202/Y204) (Cell Signaling Technology, 4370, 1:2000), mouse anti-ERK1/2 (Cell Signaling Technology, 4696, 1:2000) and mouse HRP-tagged anti-beta-actin (Santa Cruz technology, sc-47778, 1:4000). Following primary antibody incubation, blots were washed (3X) in TBST for a total of 30 min and incubated with fluorescently labelled antibodies diluted in TBST with 5% BSA, for 1.5 hours in dark at room temperature. Secondary antibodies used were as follows: Alexa Fluor goat anti-rabbit 680 (Thermo Fisher Scientific, 1:10,000), IRDye 800 CW goat anti-mouse (LI-COR Biosciences, 1:10,000) and HRP-tagged goat anti-mouse (Cell Signaling Technology, 1:2000). Following secondary antibody incubation, blots were washed (3X) for a total of 20 min and imaged on a LI-COR Odyssey fluorescence and chemiluminescence imager.

### Statistics

In each experiment, 2 or more replicate wells were included for every treatment condition. Experiments were also repeated across different days indicated as biological replicates in the figure legends. Statistical tests used for comparisons are also indicated in the figure legends.

## Acknowledgments

This work was supported by NIH RO1-DK131432-01A1 (M.N.T.) and startup funds from the Drukier Institute and Weill Cornell Medical School (M.N.T.). We thank all members of the Teruel lab and Meyer lab for helpful discussions.

## Author Contributions

S.S. and M.N.T. designed the study; S.S. and H.E.B. performed experiments and collected and analyzed data. T.M. gave conceptual advice. M.N.T. provided supervision and funding acquisition. S.S., and M.N.T. wrote the manuscript with input from all authors.

## Competing interests

The authors declare no competing interests. Correspondence and requests for materials should be addressed to M.N.T.

---

**Supplementary Figure 1.**
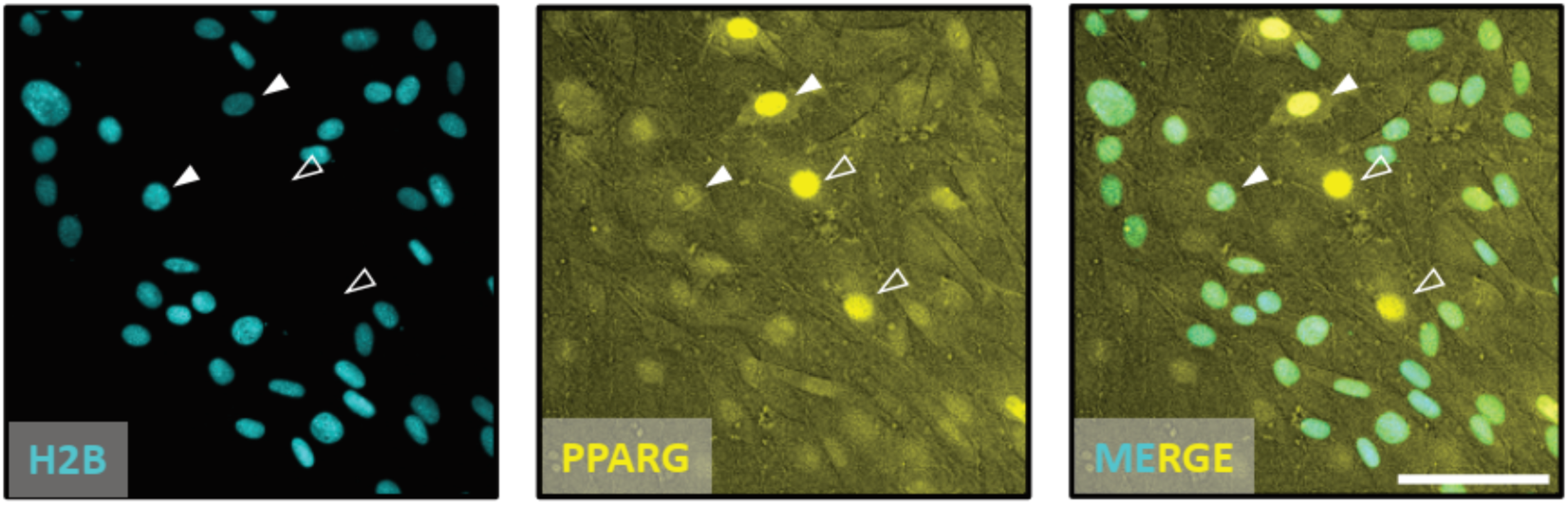
Mosaic plating for live-cell imaging and tracking of dense cultures during adipogenesis. Related to **Figure 1**. H2B-mTurquoise2 expressing cells (examples indicated by filled arrowheads) were mixed and plated with unlabeled cells (examples marked by empty arrowheads) in a 3:1 ratio for this experiment to facilitate tracking of labelled cells even as cells reaching higher densities during the 4-5 days of adipogenic differentiation protocol. Ratios of labelled to unlabelled was varied based on the changes expected from the perturbations (See Methods for more details). Scale bar - 100 µm

**Supplementary Figure 2.**
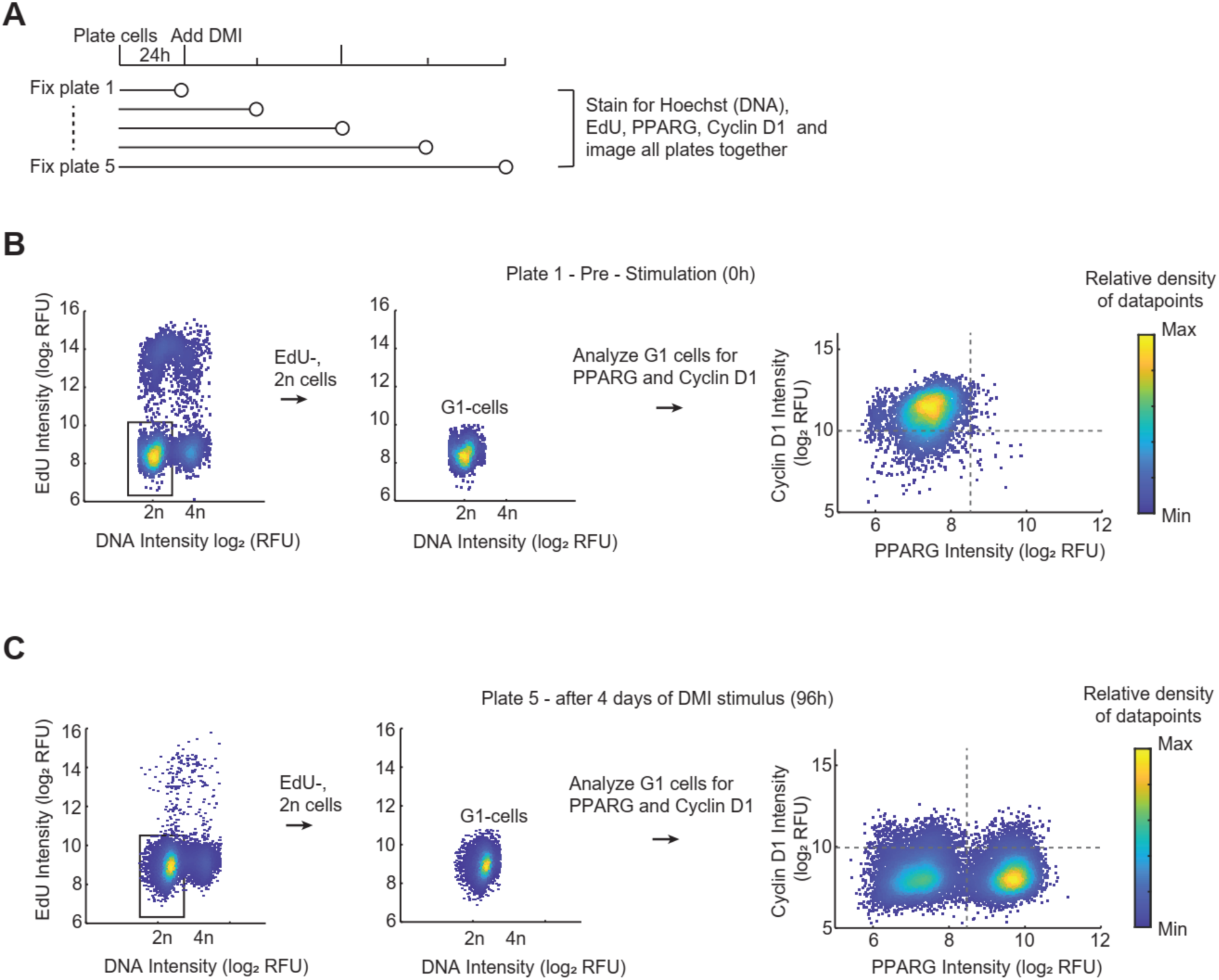
Single cell immunofluorescence analysis of Cyclin D1 levels in G1-phase cells during adipogenesis. Related to **Figure 4**. (A) Multiple 96-well plates containing with OP9 preadipocytes were subjected to the adipogenic differentiation protocol in parallel and fixed at different timepoints. All plates were then stained and imaged together. (B-C) A combination of EdU and DNA stains was used to specifically gate for G1 cells post-imaging. PPARG and Cyclin D1 levels were analyzed in such gated G1 phase cells for all the plates. Analysis pipeline for the Oh and 96h plates are shown above. Similar analyses were for performed for the measurement of p27, p21, p18 and PPARG in G1 phase cells as shown in Fig. 4B-4D.

**Supplementary Figure 3.**
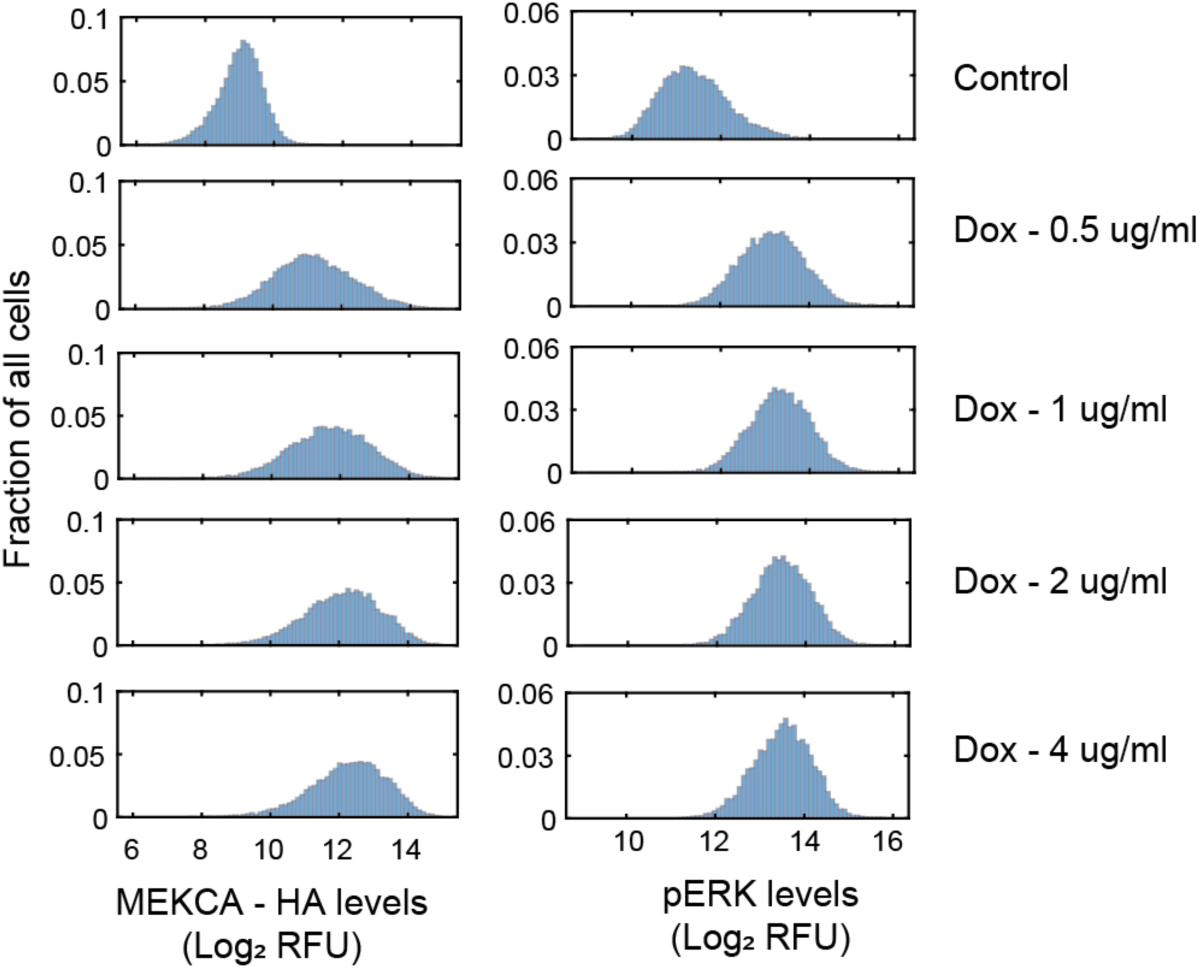
Enhancing ERK activation through expression of doxycycline-inducible constitutively active form of MEK. Related to **Figure 5**. OP9 preadipocytes were induced 24h after plating (or not in control) with increasing concentrations of doxycycline as indicated for another 24h. Cells were fixed and separate wells were stained for HA and pERK. More than 10,000 cells were analyzed from 2-3 replicate wells for each condition. Increase in the levels of MEK-CA HA expression and pERK, indicative of ERK activation, appeared to saturate at a concentration of 2 µg/ml doxycycline and hence this concentration was used for the differentiation experiments presented in Fig SH-SN.

**Supplementary Figure 4.**
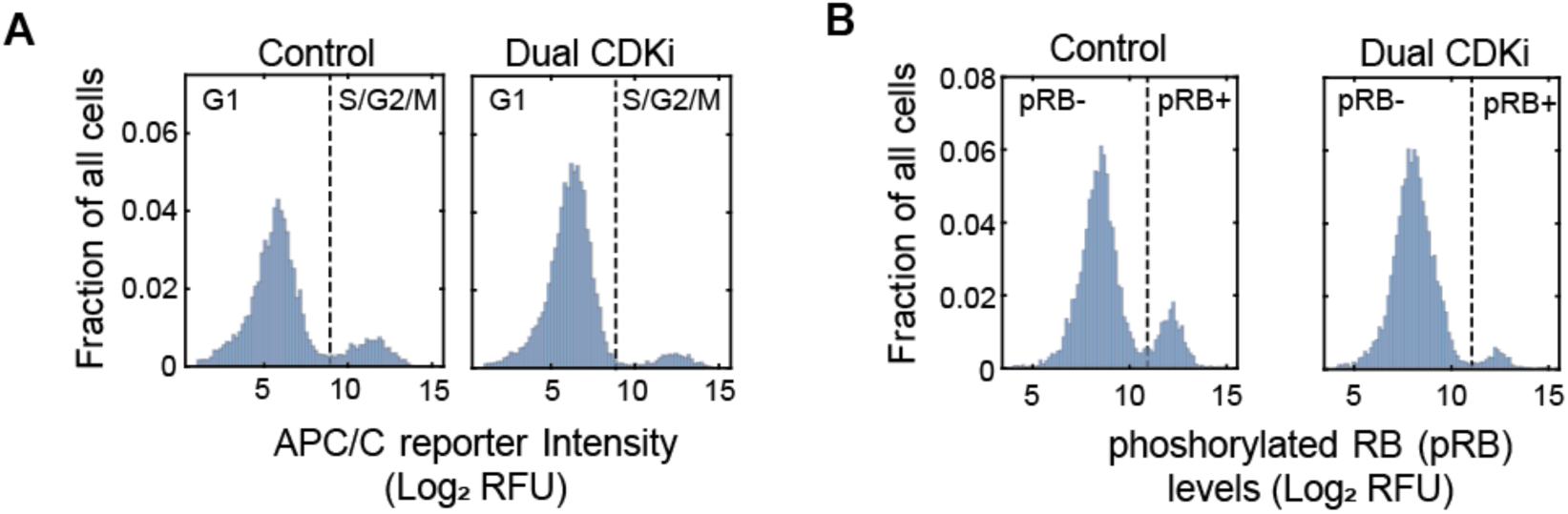
Measurement of proliferation using APC/C reporter and phosphorylation status of RB. Related to **Figure 5**. (A-8) OP9 preadipocytes expressing mCherry-APC/C reporter were fixed after 24h of treatment with 1 uM each of Palbociclib and Tagtociclib (CDK2i) or vehicle for control and stained with phospho-RB (Ser807/811) antibody. >8000 cells analyzed per condition across three replicate wells whose means are plotted in Figure SA and Figure 58. Representative of two biological replicates. (A) Histograms showing the distribution of nuclear APC/C reporter fluorescence intensities. from cells. Dashed line indicates the threshold used to identify the cells undergoing proliferation i.e. cells in S/G2/M phases with high nuclear APC/C reporter fluorescence. Also see Figure 1C. (B) Histograms showing the distribution of phosphorylated RB (pRB) antibody staining intensities from cells. Dashed line indicates the threshold used to identify the pRB+ cells i.e. cells that have entered S-phase once CDK activity is high enough to hyperphosphorylate RB.

## Notes

### Competing Interest Statement

The authors have declared no competing interest.

